# Reason’s Enemy Is Not Emotion: Engagement of Cognitive Control Networks Explains Biases in Gain/Loss Framing

**DOI:** 10.1101/109819

**Authors:** Rosa Li, David V. Smith, John A. Clithero, Vinod Venkatraman, R. McKell Carter, Scott A. Huettel

**Author notes:** indicates co-first authorship (Rosa Li, David V. Smith). Corresponding author, Box 90999, Durham, NC 27708.

## Abstract

In the classic gain/loss framing effect, describing a gamble as a potential gain or loss biases people to make risk-averse or risk-seeking decisions, respectively. The canonical explanation for this effect is that frames differentially modulate emotional processes – which in turn leads to irrational choice behavior. Here, we evaluate the source of framing biases by integrating functional magnetic resonance imaging (fMRI) data from 143 human participants performing a gain/loss framing task with meta-analytic data from over 8000 neuroimaging studies. We found that activation during choices consistent with the framing effect were most correlated with activation associated with the resting or default brain, while activation during choices inconsistent with the framing effect most correlated with the task-engaged brain. Our findings argue against the common interpretation of gain/loss framing as a competition between emotion and control. Instead, our study indicates that this effect results from differential cognitive engagement across decision frames.

**Significance Statement:** The biases frequently exhibited by human decision-makers have often been attributed to the presence of emotion. Using a large fMRI sample and analysis of whole-brain networks defined with the meta-analytic tool Neurosynth, we find that neural activity during frame-biased decisions are more significantly associated with default behaviors (and the absence of executive control) than with emotion. These findings point to a role for neuroscience in shaping longstanding psychological theories in decision science.

## Introduction

Psychologists have long described human experience as two dueling modes of thought: one process of quick emotion-laden association and another of reasoned analysis (James, 1890). More recently, a strand of research in behavioral economics has adopted a dual-process approach that contrasts automatic and low-effort Type 1 decisions against analytic and effortful Type 2 decisions (Kahneman, 2011). Efforts to identify the neural signatures of Type 1 and Type 2 decision-making (Greene et al., 2001; Sanfey et al., 2003; McClure et al., 2004; Sokol-Hessner et al., 2012) have often focused on the framing effect, in which altering how a decision is described (or “framed”) leads to systematic biases in choice (Gonzalez et al., 2005; De Martino et al., 2006; Roiser et al., 2009; Wright et al., 2012, 2013).

The canonical example of the framing effect is gain/loss framing, in which people are typically risk-averse when financial outcomes are presented as gains but risk-seeking when equivalent outcomes are presented as losses (Tversky and Kahneman, 1981). Previous studies of the neural basis of the framing effect (De Martino et al., 2006; Roiser et al., 2009; Xu et al., 2013) found amygdala activity during frame-consistent choices (i.e., risk-seeking for losses and risk-avoidance for gains) and dorsal anterior cingulate cortex (dACC) activity during frame-inconsistent choices (i.e., risk-avoidance for losses and risk-seeking for gains). Because the amygdala has been historically associated with fear and anxiety (Davis, 1992), and dACC is often associated with effortful control and conflict monitoring (MacDonald et al., 2000; Botvinick, 2007), these results have been seen as evidence for a rapid emotional brain response (Type 1) that can be overridden by effortful control (Type 2). Under this-perspective, the behavioral inconsistencies observed in the framing effect result from an intrusive emotional bias (De Martino et al., 2006; Roiser et al., 2009; Xu et al., 2013).

Yet, other evidence points to an alternative to the standard “reason vs. emotion” model, namely that framing effect arises when people adopt behavioral strategies that involve low cognitive effort. Simon’s 1955 treatise noted that “limits on computational capacity” may be the main constraint imposed upon human decision makers (Simon, 1955), and heuristic-based models of decision making emphasize that heuristics save cognitive effort (Gigerenzer and Gaissmaier, 2011; Mega et al., 2015). Emotional processing does not enter such models; decisions biases can arise, it is argued, in the absence of any emotional response. Collectively, such work provides an intriguing potential counterpoint to the standard dual-process view of the framing effect: frame-biased choices are best characterized not by high emotion but by low cognitive engagement.

In this study, we distinguished the emotion-and engagement-based explanations of the framing effect by integrating large-sample empirical functional neuroimaging data with independent maps of brain networks derived from meta-analytic tools. We analyzed functional magnetic resonance imaging (fMRI) data from a final sample of 143 participants who performed a risky decision-making task that evoked a behavioral framing effect. The resulting neural activation during frame-consistent and frame-inconsistent choices was then compared to independent meta-analytic maps from the Neurosynth database (Yarkoni et al., 2011; http://www.neurosynth.org). We found that neural activation during frame-consistent choices does include the amygdala but in fact better matches a network associated with the resting or default brain; conversely, neural activation during frame-inconsistent choices best matches a task-engaged neural network. Furthermore, we found that trial-by-trial neural similarity to resting or default networks significantly predicted frame-consistent choices, whereas trial-by-trial neural similarity to emotion-related neural networks did not. Thus, both Type 1 decisions and Type 2 decisions in gain/loss framing are best characterized along a continuum of engagement, such that Type 1 decisions reflect relative disengagement compared to Type 2 decisions.

## Materials & Methods

### Participants

Our analysis sample consisted of 143 participants (mean age 21.9 years; range 18 to 31 years; 78 female) with normal or corrected-to-normal vision and no prior history of psychiatric or neurological illness. These participants were drawn from a larger sample of 232 participants (see *Inclusion criteria* below). All participants gave written informed consent as part of a protocol approved by the Institutional Review Board of Duke University Medical Center. We note that this is much larger than most imaging studies, allowing us ample statistical power to detect effects (Button et al., 2013).

### Stimuli and task

Participants performed 126 trials of a risky decision-making task (Figure 1) adapted from previous studies of the neural basis of framing effects (De Martino et al., 2006), split across three runs by short breaks. On each trial, participants were shown a starting amount that varied uniformly from $8 to $42. Participants were then asked to choose between “safe” and “gamble” options with a button press. Left and right positions of the safe and gamble options were randomized across trials. Safe options were framed such that participants could keep (gain frame) or lose (loss frame) a subset of the starting amount for sure. The gamble option did not differ according to frame and was represented by a pie chart reflecting probabilities to “Keep All” or “Lose All” of the original starting amount (20%-80%, 25%-75%, 33%-67%, 50%-50%, 67%-33%, or 75%-25%). The expected value varied between the safe and gamble options (safe: gamble EV ranged from 0.24 to 3.08). On half of the trials, participants played for themselves, and on the other half of the trials, they played for a charity of their choice (Animal Protection Society of Durham, Durham Literacy Center, Easter Seals UCP North Carolina, or American Red Cross: The Central North Carolina Chapter). Nearly all trials (92%) were matched with an identical trial in the opposite frame.

**Figure 1.**
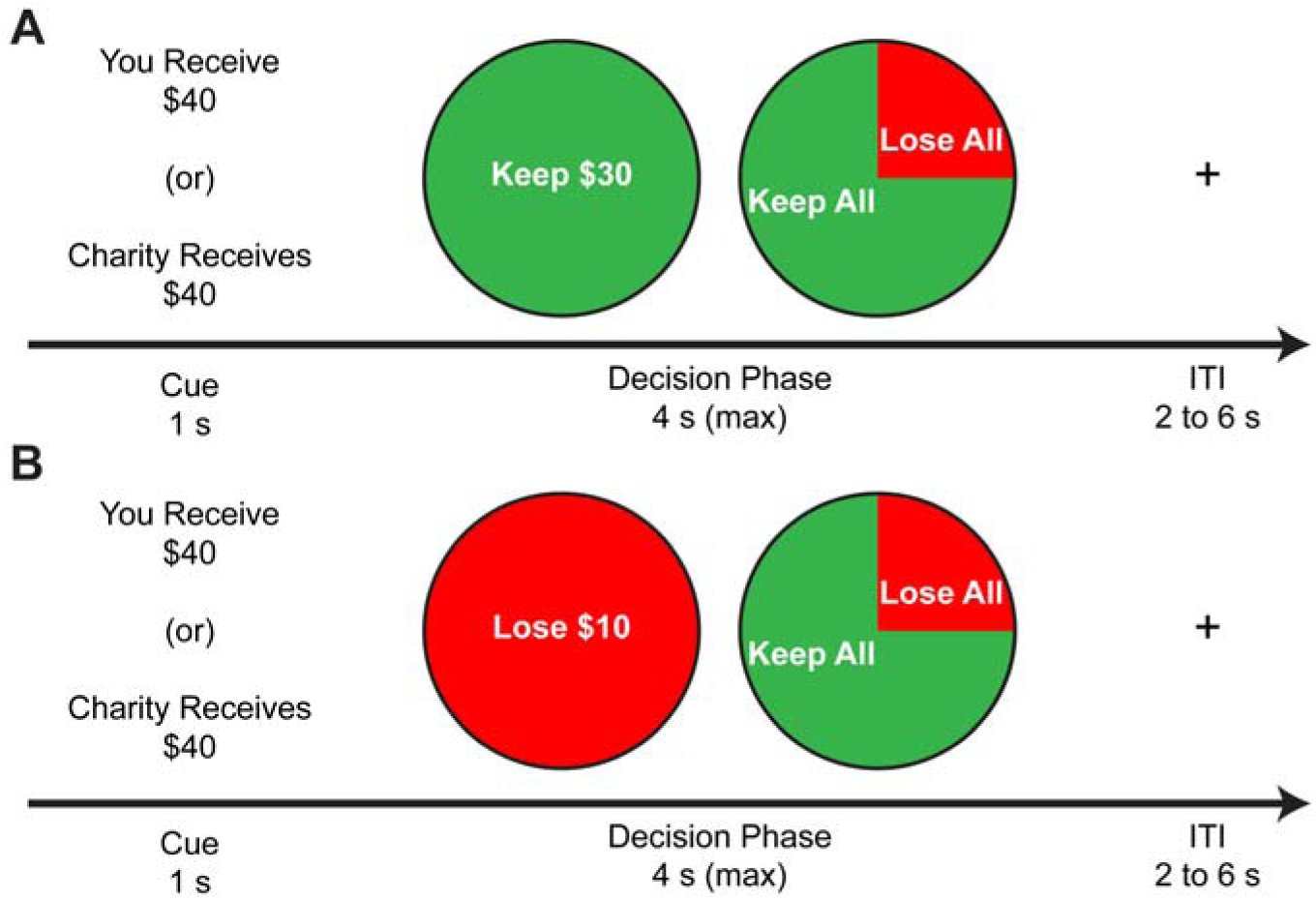
Participants engaged in a financial decision-making task.At the beginning of each trial, an initial endowment indicated the target of the decision (self or charity). Following this cue, participants had the opportunity to choose between an all-or-nothing gamble or a safe option with a guaranteed proportion of the initial endowment. The safe option was presented in two conditions: **(A)** a gain frame; and **(B)** a loss frame. Crucially, nearly all trials were matched to an identical trial in the opposite frame that only differed in the presentation of the safe option (‘keep’ or ‘lose’). After the choice, a fixation cross was presented for 2 to 6 seconds. At the end of the experiment, one trial chosen at random was resolved for payment.

Stimuli were projected onto a screen at the back of the scanner bore, and participants viewed the stimuli through mirrored goggles. Tasks were programmed using the Psychophysics Toolbox version 2.54 (Brainard, 1997). At the end of the experiment, one trial was randomly selected to be carried out for potential payment in order to ensure incentive compatibility across all trials. Participants also completed two additional fMRI tasks and a resting state scan (described in Utevsky et al., 2014) that were not analyzed for this manuscript.

### Behavioral analysis

The behavioral framing effect was calculated using a standard metric: the difference between the percentage of gamble choices in the loss frame relative to the percentage of gamble choices in the gain frame (De Martino et al., 2006). Trials in which no choice was made were excluded from the calculation of this metric.

For response time analyses, choices were classified as either frame-consistent or frame-inconsistent. Frame-consistent choices were safe decisions in the gain frame (Gain_safe_) and gamble decisions in the loss frame (Loss_gamble_). Frame-inconsistent choices were gamble decisions in the gain frame (Gain_gamble_) and safe decisions in the loss frame (Loss_safe_).

### Image acquisition

Functional MRI data were collected using a General Electric MR750 3.0 Tesla scanner equipped with an 8-channel parallel imaging system. Images sensitive to blood-oxygenation-level-dependent (BOLD) contrast were acquired using a T_2_*-weighted spiral-in sensitivity encoding sequence (acceleration factor = 2), with slices parallel to the axial plane connecting the anterior and posterior commissures [repetition time (TR): 1580 ms; echo time (TE): 30 ms; matrix: 64 x 64; field of view (FOV): 243 mm; voxel size: 3.8 x 3.8 x 3.8 mm; 37 axial slices acquired in an ascending interleaved fashion; flip angle: 70°]. We chose these sequences to ameliorate susceptibility artifacts in ventral frontal regions (Pruessmann et al., 2001; Truong and Song, 2008). Prior to preprocessing these functional data, we discarded the first eight volumes of each run to allow for magnetic stabilization. To facilitate coregistration and normalization of these functional data, we also acquired whole-brain high-resolution anatomical scans (T_1_-weighted FSPGR sequence; TR: 7.58 ms; TE: 2.93 ms; voxel size: 1 x 1 x 1 mm; matrix: 256 x 256; FOV: 256 mm; 206 axial slices; flip angle: 12°).

### Preprocessing

Our preprocessing employed tools from the FMRIB Software Library (FSL Version 4.1.8; http://www.fmrib.ox.ac.uk/fsl/; RRID: SCR_002823) package (Smith et al., 2004; Woolrich et al., 2009). We first corrected for head motion by realigning the time series to the middle time point (Jenkinson et al., 2002). We then removed non-brain material using the brain extraction tool (Smith, 2002). Next, intravolume slice-timing differences were corrected using Fourier-space phase shifting, aligning to the middle slice (Sladky et al., 2011). Images were then spatially smoothed with a 6 mm full-width-half-maximum Gaussian kernel. To remove low-frequency drift in the temporal signal, we then subjected the functional data to a high-pass temporal filter with a 150 second cutoff (Gaussian-weighted least-squares straight line fitting, with sigma = 50 s). Finally, each 4-dimensional dataset was grand-mean intensity normalized using a single multiplicative factor. Prior to group analyses, functional data were spatially normalized to the MNI avg152 T1-weighted template (2 mm isotropic resolution) using a 12-parameter affine transformation implemented in FLIRT (Jenkinson and Smith, 2001); these transformations were later applied to the statistical images before cross-run and cross-participant analyses.

### fMRI analysis

Neuroimaging analyses were conducted using FEAT (FMRI Expert Analysis Tool) Version 5.98 (Smith et al., 2004; Woolrich et al., 2009). Our first-level analyses (i.e., within-run) utilized a general linear model with local autocorrelation correction (Woolrich et al., 2001) consisting of four regressors modeling each frame and participant choice (Gain_safe_, Gain_gamble_, Loss_safe_, Loss_gamble_). We defined the duration of each regressor as the period of time from the presentation of the cue to the time of choice. This procedure controls for confounding effects related to response time (Grinband et al., 2011a, 2011b). In this first-level model, we also included two 0-second impulse regressors to account for the presence of the initial cue and for the value of that cue, and we included nuisance regressors to account for missed responses, head motion, and outlier volumes. Except for the head motion and outlier volume nuisance regressors, all regressors were convolved with a canonical hemodynamic response function. We combined data across runs, for each participant, using a fixed-effects model, and combined data across participants using a mixed-effects model (Beckmann et al., 2003; Woolrich et al., 2004).

Our group-level analyses included two contrasts of interest (De Martino et al., 2006; Roiser et al., 2009; Xu et al., 2013). Neural activity associated with the framing effect was modeled by contrasting frame-consistent choices over frame-inconsistent choices (see Equation 1a). Frame-inconsistent neural activity was modeled by the reversed contrast (frame-inconsistent choices contrasted over frame-consistent choices; see Equation 1b).

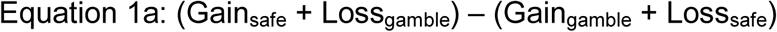

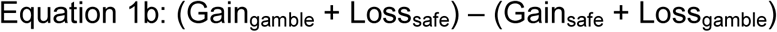

Statistical significance was assessed using Monte Carlo permutation-based statistical testing with 10,000 permutations (Nichols and Holmes, 2002; Winkler et al., 2014). Additionally, we used threshold-free cluster enhancement to estimate clusters of activation that survived a corrected family-wise-error-rate of 5% (Smith and Nichols, 2009). Statistical overlay images were created using MRIcron and MRIcroGL (Rorden and Brett, 2000). All coordinates are reported in MNI space.

### Inclusion criteria

Given our sample size, we adopted stringent *a priori* criteria for data quality to determine inclusion/exclusion of participants. First, we estimated the average signal-to-fluctuation-noise ratio (SFNR) for each run (Friedman et al., 2006). Second, we computed the average volume-to-volume motion for each run. Third, we identified outlier volumes in our functional data. We considered a volume an outlier if its root-mean-square (RMS) amplitude exceeded the value of 150% of the interquartile range of RMS for all volumes in a run. Using these three metrics, we excluded runs in which any measure metric was extreme relative to the other runs (i.e., SFNR < 5^th^ percentile of the distribution of SFNR values; outlier volumes > 95^th^ percentile the distribution of outlier volumes; average volume-to-volume motion > 95^th^ percentile). Runs with an excessive number of missed behavioral responses (>97.5^th^ percentile all runs, or >26.2% of trials with no responses) were also excluded from analyses. Data from two additional participants were excluded due to poor registration to the template brain. This resulted in the exclusion of 28 participants.

After excluding runs based on the above criteria, our final analyses excluded all participants who had fewer than two runs that each had at least two trials of each regressor type (i.e.: Gain_safe_, Loss_gamble_, Gain_gamble_, Loss_safe_). As examples, participants who always chose the safe option or always chose the gamble option were not included in the analyses, since no contrast could be constructed between their choices. This resulted in the exclusion of an additional 61 participants, resulting in a final model that included 143 participants, each with multiple runs of high-quality data and behavior mixed between frame-consistent and frame-inconsistent choices.

### Neural similarity analysis

We used the fMRI meta-analysis software package Neurosynth (Yarkoni et al., 2011; version 0.3; RRID: SCR_006798) to construct metrics of neural similarity between our empirical fMRI data and reverse-inference maps drawn from prior studies (i.e., the 8,000+ published, peer-reviewed studies included in the Neurosynth database at the time of our analyses).

We calculated Pearson correlation coefficients between the unthresholded Z-statistic map of our framing effect contrast (referred to as Framing Contrast; see Equation 1a) and each of Neurosynth’s 2,592 term-based reverse-inference *Z*-statistic maps (referred to as Neural Profiles) for all voxels within our group-level brain mask using the Neurosynth Decode tool (Yarkoni et al., 2011).

Using the Pearson correlation coefficients between our Framing Contrast and the Neurosynth term-based maps, we identified the 10 most positively correlated and 10 most negatively correlated Neural Profiles (referred to as NP+ and NP−, respectively, and NP± when referring to both sets of maps), as well as 6 emotion-related Neural Profile (the Neurosynth maps for “emotions”, “feelings”, “emotion”, “emotionally”, “amygdala”, and “feeling”; hereafter referred to as NPe). We then used partial correlation analyses to compare the shared variance between our empirical results and the maps drawn from Neurosynth (Figure 2).

**Figure 2.**
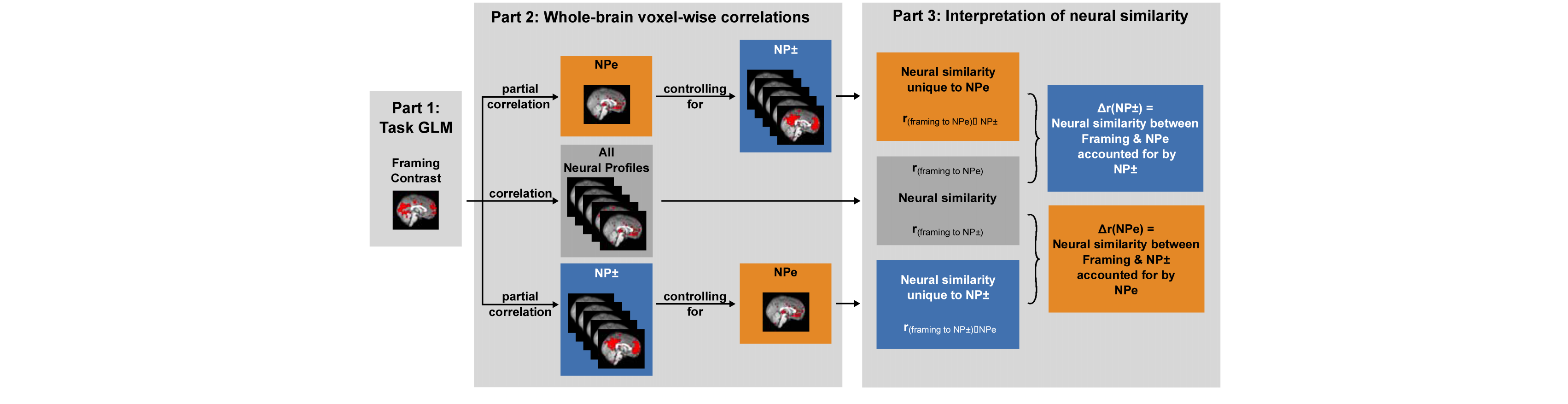
Neural similarity measures were obtained by correlating our whole-brain Framing Contrast with Neurosynth’s reverse-inference term-based maps, or Neural Profiles (NP). Unique neural similarity was calculated using partial correlation analyses for a specific term while controlling for additional terms. The reduction in neural similarity following such partial correlation analyses was attributed to the controlling terms. See *Experimental Procedures: Neural similarity analysis* for details.

For each NP±, we quantified its unique neural similarity to the Framing Contrast as its partial correlation coefficient with our Framing Contrast after controlling for the NPe, or r_(framing to NP±)•NPe_. We then subtracted this partial correlation coefficient from the correlation coefficient between our Framing Contrast and each NP± (r_(framing to NP±)_–r_(framing to NP±)•NPe_). This absolute difference, orr(NPe), is the neural similarity between our Framing Contrast and each NP± that is accounted for by each NPe.

Similarly, for each NPe, we quantified its unique neural similarity to the Framing Contrast as its partial correlation coefficient with our Framing Contrast map after controlling for each NP±, or r_(framing to NPe)•NP±_. We then subtracted each partial correlation coefficient from the correlation coefficient between our Framing Contrast and each NPe (r_(framing to NPe)_ – r_(framing to NPe)•NP±_). This absolute difference, or Δr(NP±), is the neural similarity between our Framing Contrast and each NPe that is accounted for by each NP±.

### Trial-by-trial analysis

We estimated an additional first-level trial-by-trial general linear model with a separate regressor for each trial (42 trials per run) and the same impulse and nuisance regressors of our Framing Contrast first-level model (excepting the missed response trial nuisance regressors). This approach allowed us to capture variance tied to each individual trial and characterize trial-specific processes (e.g., frame-consistent or - inconsistent choices). For each trial, we obtained a whole-brain z-score map. These trial-level z-score maps were correlated with each NP± and NPe map to derive trial-by-trial fluctuations in neural similarity. We averaged the Pearson correlation coefficients for terms within each NP+, NP−, and NPe group, thus yielding a single neural similarity score for each NP group.

We then used these trial-level neural similarity scores as regressors in two logistic regression models predicting frame-consistent decisions. Our Basic Model used response time and neural similarity scores for the NP+, NP−, and NPe as regressors. Our Interaction Model used response time, neural similarity scores for the NP+, NP−, and NPe, and the interaction of mean-centered NP+ and NPe as regressors. Thus, for each participant, we obtained regression coefficients summarizing the degree to which neural similarity to the NP+, the NP−, the NPe, the interaction of NP+ and NPe, and response time predicted frame-consistent decisions at the trial-by-trial level.

### Data Availability

Our group-level unthresholded Framing Contrast is deposited in Neurovault (Gorgolewski et al., 2015)http://neurovault.org/collections/ECNKKIIS/

## Results

### Decision frames alter response time and gambling behavior

We examined response times as functions of decision target (self and charity) and frame type (gain and loss) and as functions of decision target and choice (frame-consistent and frame-inconsistent) using 2x2 repeated measures ANOVAs. We found that participants were slower to respond in loss frames (mean = 1.87s, SEM = 0.04s) compared to gain frames (mean = 1.72s, SEM = 0.03s; *F*
_(1,142)_ = 175.55, *p* < 0.001; Figure 3A). Participants were also slower to make frame-inconsistent choices (Gain_gamble_ and Loss_safe_; mean = 1.92s, SEM = 0.04s) than frame-consistent choices (Gain_safe_ and Loss_gamble_; mean = 1.75s, SEM = 0.03s; *F*
_(1,142)_ = 180.54, *p* < 0.001). In contrast, decision target did not significantly modulate response times (self mean = 1.81s, SEM = 0.03s; charity mean = 1.78s, SEM = 0.04s; *F*
_(1,142)_ = 3.40, *p* = 0.0673). Our analyses also indicated that decision target did not significantly interact with frame type (*F*(_1,142)_ = 2.63, *p* = 0.1072).

Participants exhibited a behavioral framing effect and increased their gambling behavior in loss frames (mean = 54.98%, SEM = 1.60%) relative to gain frames (mean=36.62%, SEM = 1.35%; *F*
_(1,142)_ = 180.95, *p* < 0.001; Figure 3B). Though the task was originally designed so that we could evaluate potential differences in the framing effect depending on the target of decisions (i.e. self vs. charity), participants showed no differences in overall gambling behavior between the self (mean = 45.56%, SEM =1.42%) and charity (mean = 46.04%, SEM = 1.40%) decision targets (*F*_(1,142)_ = 0.21, *p* =0.6454), nor was there a target-by-frame interaction (*F*_(1,142)_ = 0.30, *p* = 0.5820). In addition, although participants varied substantially in their responses to the framing manipulation, we found that the extent to which the framing manipulation biased a participant’s choices was largely consistent across self and charity decision targets (*r*_(141)_ = 0.59, *p* < 0.001; Figure 3C). Taken together, these results confirm that our participants exhibited the hallmarks of a framing effect – increased risk aversion for the gain frame and slower decisions for frame-inconsistent choices – and that framing effects were consistent across the two decision targets. Because our behavioral analyses indicated that framing effect did not depend on decision target, we collapsed across decision target in our neuroimaging analyses. This provides increased power by doubling the number of trials in each condition.

**Figure 3:**
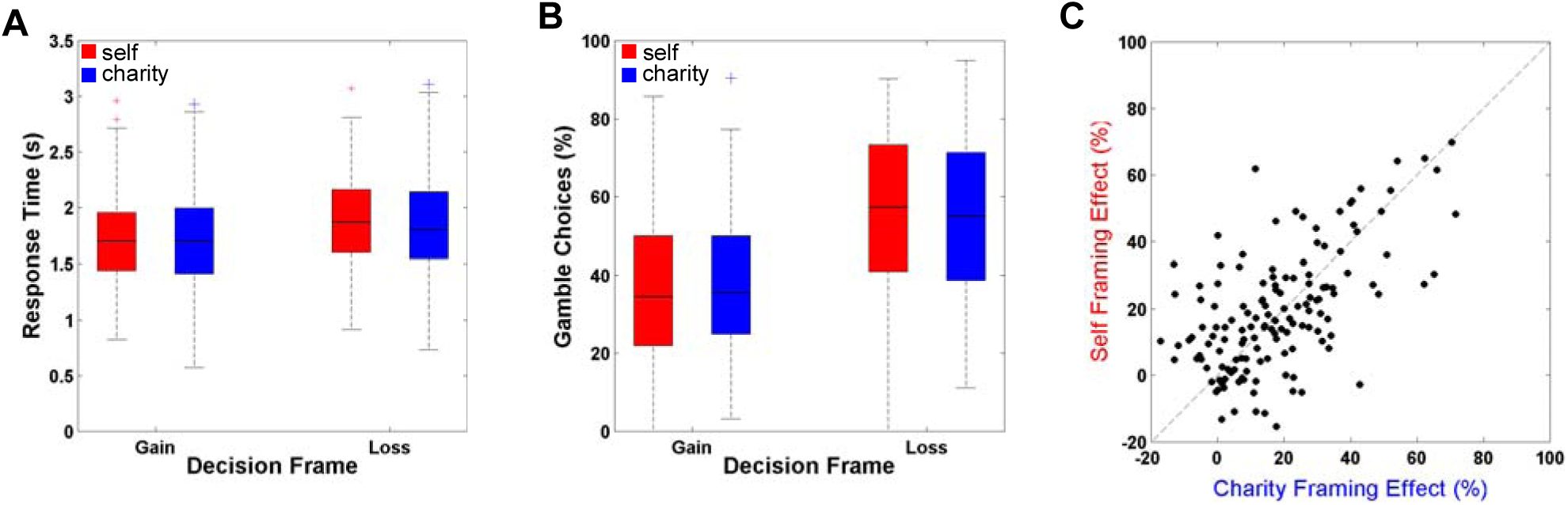
Decision frame (loss or gain) but not decision target (charity or self) significantly affected behavioral response time and choice (n = 143 participants). (A) Participants responded more slowly during the loss frame compared to the gain frame. This effect did not depend on whether the outcome of the decision was intended for the participant or the participant’s charity. Standard boxplots are shown. (B) When decisions were framed as a potential loss (relative to a potential gain), participants gambled more, indicating a robust framing effect. Notably, the magnitude of the framing effect -- indexed by the proportion of gambles in the loss frame relative to the gain frame -- did not depend on whether the outcome of the decision was intended for the participant or the participant’s charity. Standard boxplots are shown. (C) Individual differences in the magnitude of the framing effect (% loss_gamble_ – % gain_gamble_) were consistent within participants for self and charity targets.

### Decision frames change brain networks associated with cognitive engagement

Our Framing Contrast (frame-consistent choices > frame-inconsistent choices) found activation during frame-consistent choices that replicated the significant amygdala activation found in previous studies (De Martino et al., 2006), some of which used region-of-interest analyses (Roiser et al., 2009; Xu et al., 2013). In our model, this amygdala activation was bilateral and significant after whole-brain correction (MNI coordinates: [26, 0, -22] and [-22, 6, 18];
Figure 4A). We also found, however, significant and widespread activation during frame-consistent choices throughout the brain after whole-brain correction, including but not limited to bilateral insula, posterior cingulate cortex (PCC), and dorsal medial prefrontal cortex (dmPFC; Figure 5A).

**Figure 4.**
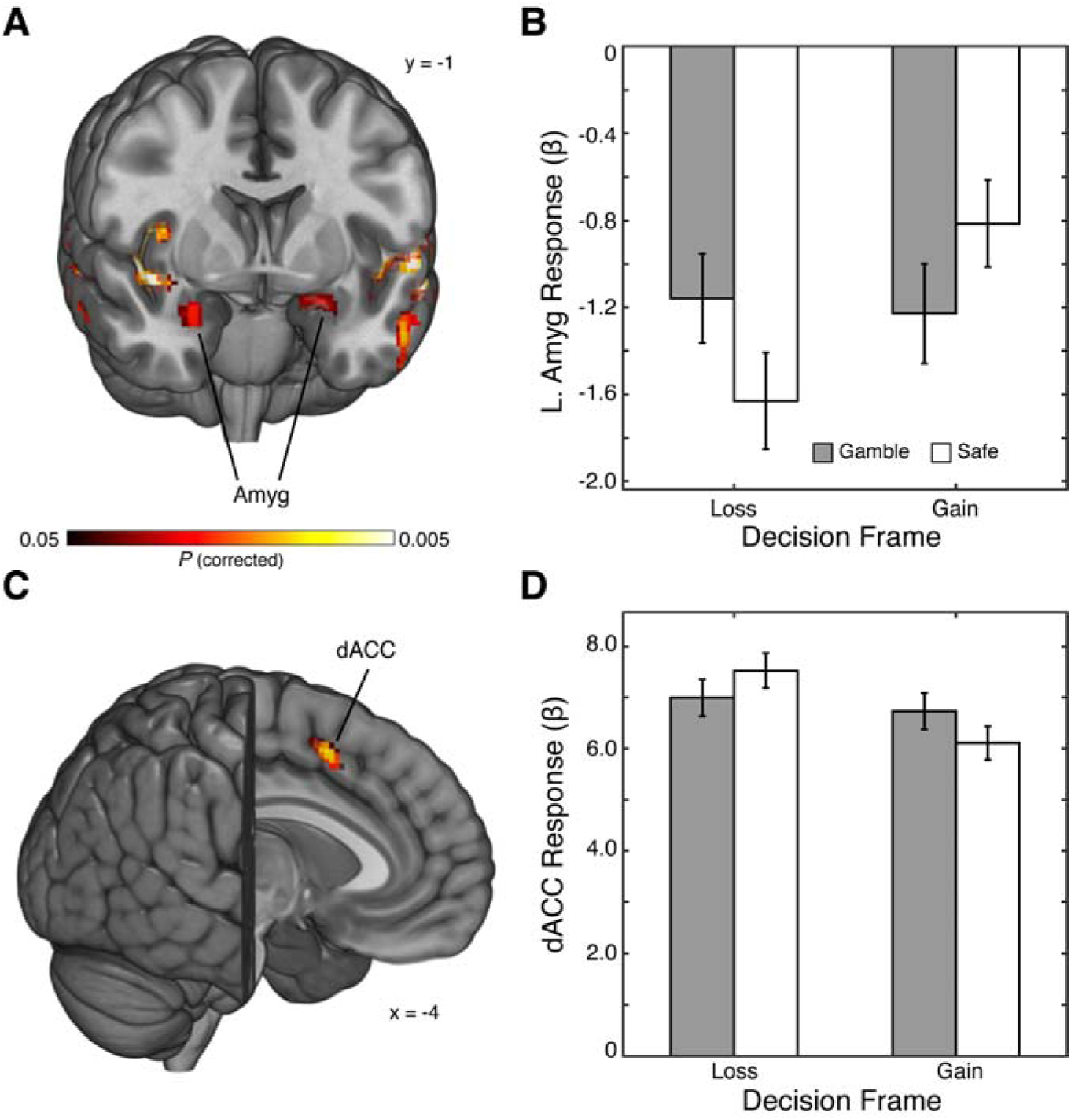
We used an interaction contrast (see*Experimental Procedures: Equation 1a*) to identify regions with greater activation toframe-consistent choices than to frame-inconsistent choices. This analysis revealed widespread activation, including bilateral amygdala. **(B)** Within the amygdala, we found increased response to gamble choices relative to safe choices when the decision was framed as a potential loss. In contrast, when the decision was framed as a potential gain, we observed increased responses to safe choices relative to gamble choices. **(C)** We used an interaction contrast (see *Experimental Procedures*: Equation 1b) to identify regions with greater activation to frame-inconsistent choices than to frame-consistent choices. This analysis revealed activation within dorsal anterior cingulate cortex (dACC). **(D)** Within dACC, we found increased response to safe choices relative to gamble choices when the decision was framed as a potential loss. In contrast, when the decision was framed as a potential gain, we observed increased responses to gamble choices relative to safe choices. Reported activations are shown at p < 0.05 (corrected 2-tailed t-test). Error bars reflect SEM.

Using the reverse of the Framing Contrast (frame-inconsistent choices > frame-consistent choices) to identify neural activity associated with frame-inconsistent choices, we replicated previous findings of significant activation in dACC (De Martino et al., 2006; Roiser et al., 2009; Murch and Krawczyk, 2013; Xu et al., 2013; MNI coordinates: [-4, 12, 46]; Figure 4C
). We also found significant activation in left inferior frontal gyrus during frame inconsistent choices (Figure 5A).

Our neural similarity analyses found that our Framing Contrast was more correlated with Neural Profiles associated with the default or resting brain (e.g. state, default, resting, mode, PCC) than with Neural Profiles associated with emotion (e.g. emotions, feeling, amygdala). Conversely, the reverse of the Framing Contrast was most correlated with Neural Profiles associated with effortful processing (e.g. working, task, executive, frontal, maintenance, load; Figure 5B).

**Figure 5:**
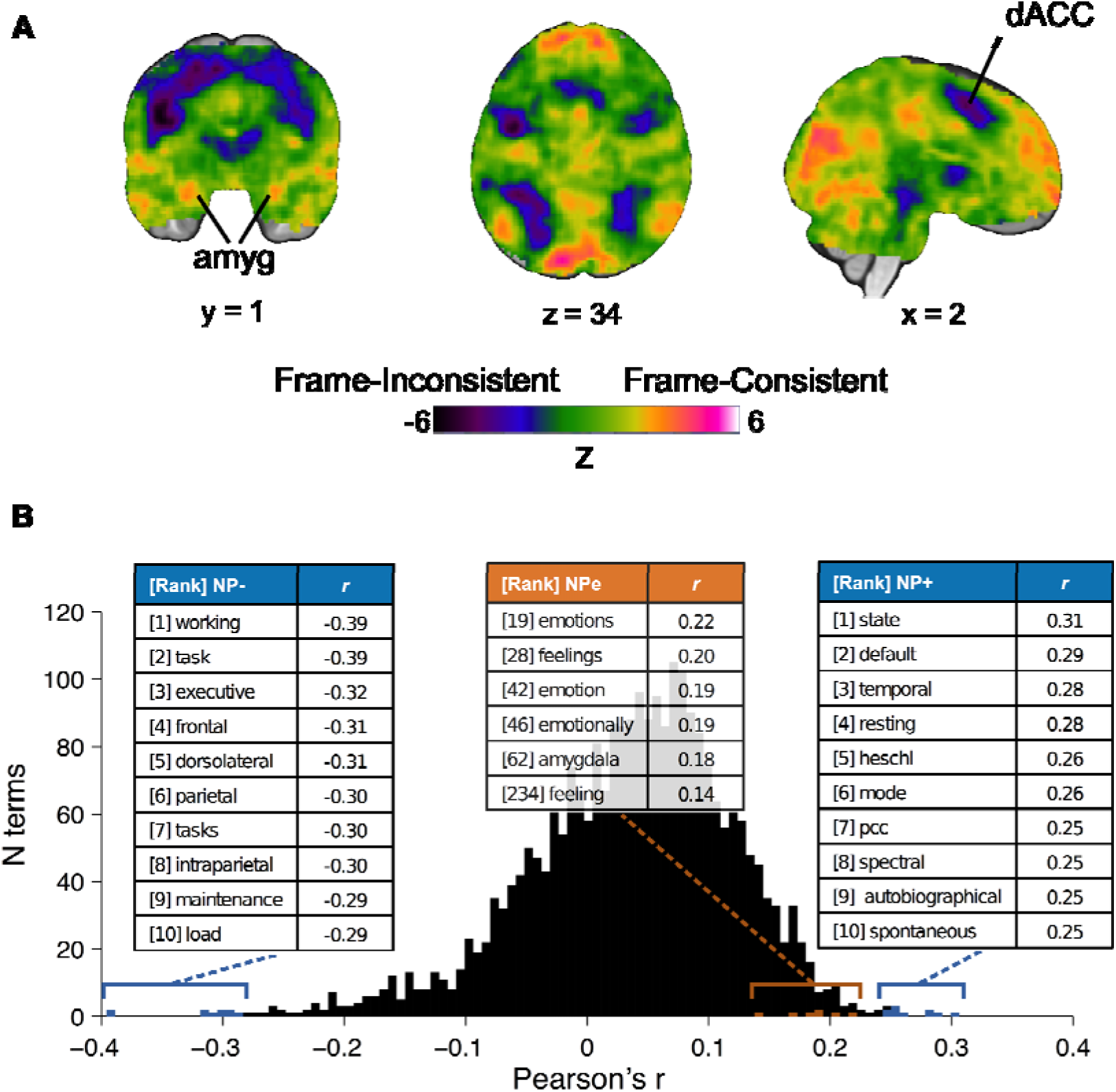
Neural activity during frame-consistent choices resembles the defaul t brain, while neural activity during frame-inconsistent choices resembles the task - engaged brain. Given the diffuse nature of our neural framing effects, we depict the unthresholded statistical images across the whole brain for both contrasts. (A) Activation associated with frame-consistent choices (hot colors; see *ExperimentalProcedures*: Equation 1a) resembled the default-mode network. In contrast, activationassociated with frame-inconsistent choices (cool colors; see *Experimental Procedures*: Equation 1b) resembled the executive control network. Images are unthresholded toshow whole-brain activity. (B) To systematically compare our activations with those of over 8,000 previously reported studies, we computed spatial correlations between the unthresholded Z-statistic map of our Framing Contrast and the reverse-inference Z-statistic maps of each of the 2,592 terms in the Neurosynth database (Neural Profiles, or NP). Pearson’s r-values for each paired correlation are shown in the histogram. The 10 most correlated (NP+) and anti-correlated (NP−) terms are highlighted in blue, and emotion-related terms are highlighted in orange.

Additionally, our partial correlation analyses found that for each NPe, the neural similarity accounted for by the NP±, or Δr(NP±), was significantly greater than the neural similarity accounted for by each NPe, or Δr(NPe) (for all NP±, mean = 0.043, SEM = 0.003; for all NPe, mean = 0.022, SEM = 0.001; 2-sided paired t-test *t*(238) = 6.323, *p* < 0.001). Thus, the variance shared by our Framing Contrast with the NPe and NP± is better accounted for by the NP± than by the NPe (Figure 6).

**Figure 6:**
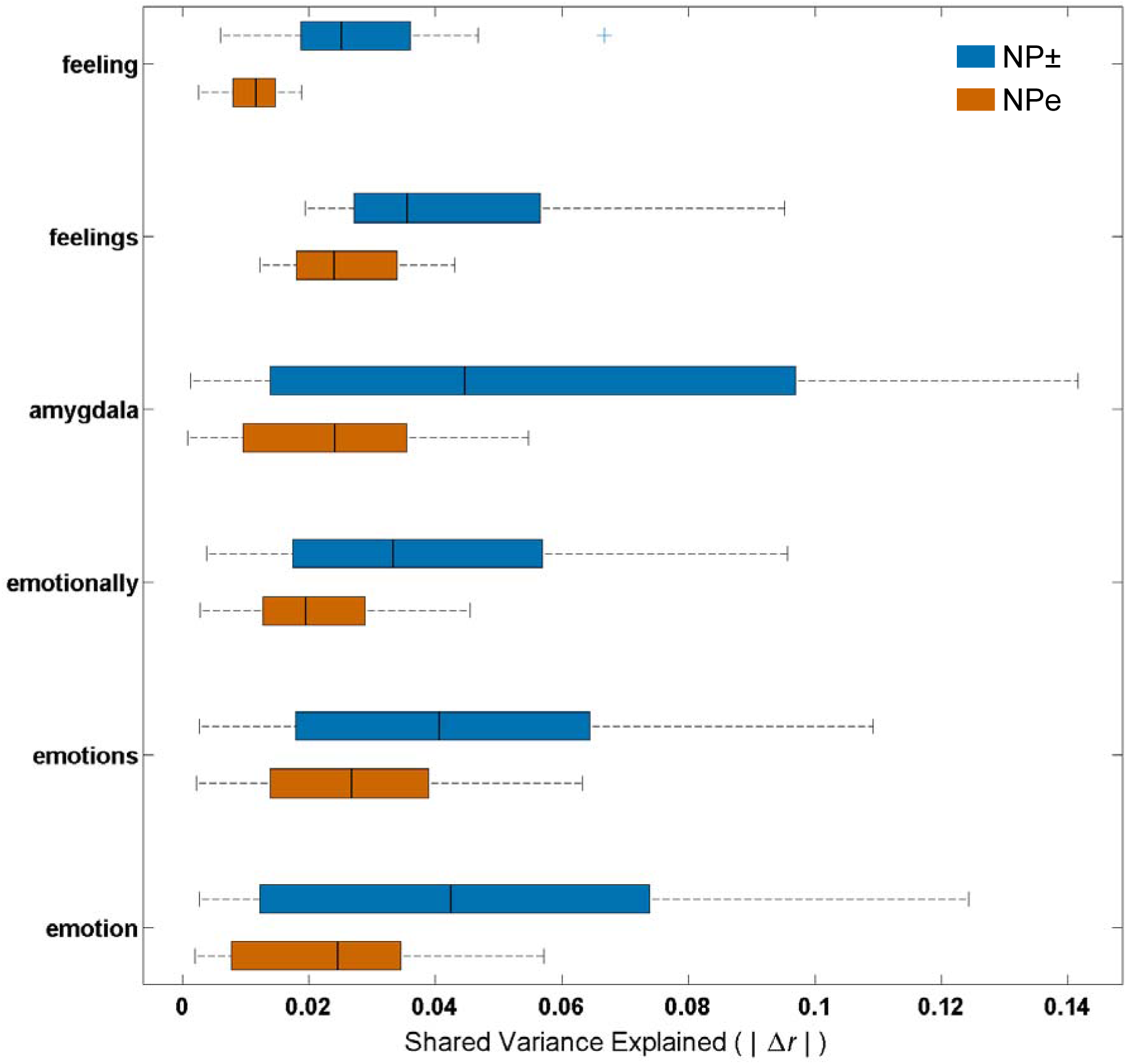
Shared variance between fMRI statistical maps for the Framing Contrast and Neurosynth term-based maps is better accounted for by Neural Profiles associated with the default and task-engaged brain (NP±) than by Neural Profiles associated with emotion-related terms (NPe). Orange bars show the neural similarity between the Framing Contrast and all NP± that is accounted for each NPe or 
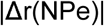
. Blue bars show the neural similarity between the Framing Contrast and each NPe that is accounted for by all NP±, or 
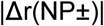
.

We additionally examined whether positive and negative emotion-related terms were differentially associated with the framing contrast map. We found that positive (or approach-related) Neural Profiles exhibited low neural similarity scores: affect (0.078), affective (0.167), anticipation (−0.061), approach (0.107), arousal (0.076), gain (−0.115), and happy (0.157). We also found that negative (or avoidance-related) Neural Profiles exhibited similarly low neural similarity scores: anger (0.172), angry (0.094), anxiety (0.065), avoid (0.098), disgust (0.172), fear (0.141), fearful (0.147), loss (0.109), and pain (0.02). [We note that other relevant terms (joy, trust, surprise) are not contained in the Neurosynth database.] These results provide further support for the claim that neither emotion nor emotional valence are robust predictors of the patterns of brain responses associated with the framing effect.

### Trial-by-trial cognitive disengagement predicts frame-consistent choices

We used trial-by-trial fluctuations in neural similarity to the NP+, NP−, and NPe to test whether trial-level neural similarity to cognitive disengagement (or cognitive engagement) predicts choices that are frame-consistent (or frame-inconsistent). Our Basic Model found that frame-consistent choices were significantly predicted by trial-level neural similarity to the NP+ (terms associated with the disengaged brain) and response time (NP+: mean beta = 2.51, SEM = 0.55, t(142) = 4.54, p < 0.001; RT: mean beta = −0.60, SEM = 0.05, t(142) = −11.09, p < 0.001) but not by trial-level neural similarity to the NP− or NPe (terms associated with the engaged brain or emotion; NP−: mean = −0.49, SEM = 0.40, t(142) = −1.21, p = 0.23; NPe: mean = −1.00, SEM = 0.62, t(142) = −1.62, p = 0.11; Figure 7
).

**Figure 7:**
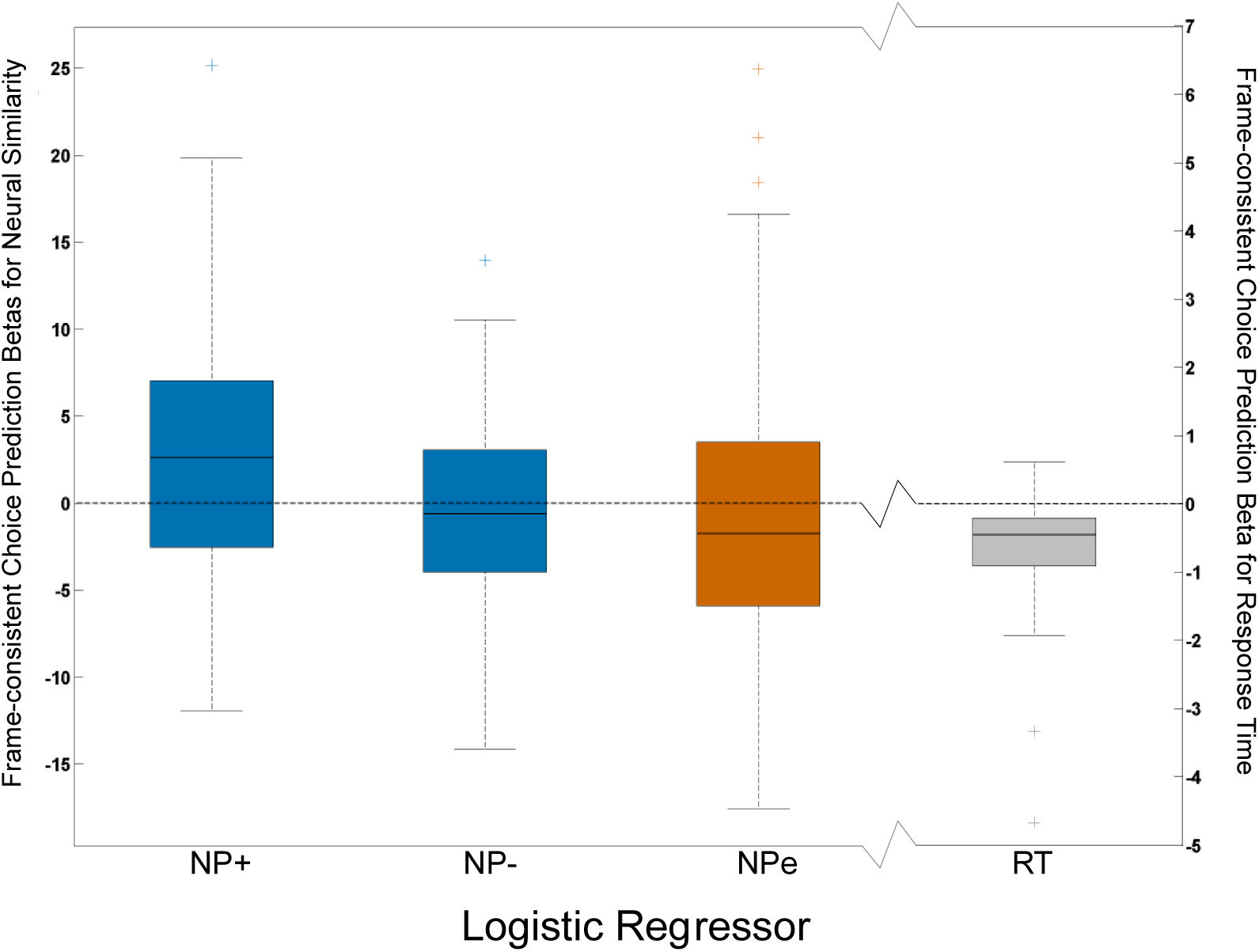
Trial-by-trial neural similarity to Neural Profiles associated with the default brain (NP+) and response time significantly predict frame-consistent choices. We used logistic regression to predict whether a subject made a frame-consistent or frame-inconsistent choice on each trial. Four regressors were entered into the model (see Methods): average similarity to Neural Profiles associated with the default brain (NP+), average similarity to Neural Profiles associated with the task-engaged brain (NP−), average similarity to Neural Profiles associated with emotion-related terms (NPe), and response time (RT). Greater neural similarity to NP+ (leftmost blue bar) and shorter response times (grey bar) significantly predicted frame-consistent choices.

To test whether the relationship between trial-level neural similarity to NP+ and NPe significantly predicted frame-consistent choice, we conducted an additional logistic regression analysis that added the interaction of NP+ and NPe to the Basic Model. In this Interaction Model, response time and trial-level neural similarity to NP+ still significantly predicted frame-consistent choices (NP+: mean beta = 2.53, SEM = 0.57, t(142) = 4.41, p < 0.001; RT: mean beta = −0.61, SEM = 0.05, t(142) = −11.24, p < 0.001), while trial-level neural similarity to NP−, NPe, and the interaction of NP+ and NPe did not (NP−: mean beta = −0.57, SEM = 0.41, t(142) = −1.40, p = 0.16; NPe: mean beta = −1.06, SEM = 0.65, t(142) = −1.64, p = 0.10; NP+*NPe: mean beta = 9.11, SEM = 5.91, t(142) = 1.54, p = 0.13), Furthermore, average AIC and BIC were significantly lower for the Basic Model (mean AIC = 149.97, mean BIC = 163.43) compared to the Interaction Model (mean AIC = 150.79, mean BIC = 166.94; AIC paired t(142) = -6.34, p < 0.001; BIC paired t(142) = -27.11, p < 0.001), indicating that the Basic Model had a better fit compared to the Interaction Model.

We further investigated whether trial-level neural similarity to cognitive disengagement was predictive of frame-consistent choices within only the subset of gain-framed trials or only the subset of loss-framed trials. Basic Model results within gain-and loss-framed subsets were qualitatively identical to the results for all trials. That is, for both gain-and loss-framed trials, frame-consistent choices were significantly predicted by trial-level neural similarity to the NP+ and response time, but not by trial-level neural similarity to the NP− or NPe (see Table 1). We note that these results indicate that neural similarity to NP+ predicts opposing choices in the different frames: in gain-framed trials, it predicts safe choices, while in loss-framed trials, it predicts gamble choices.

**Table 1:** Results of Basic Model using subsets of trials to predict frame-consistent choice from trial-level neural similarity and response time

| Trials subset | Regressor | Mean beta (SEM) | *t*-statistic | *p*-value |
| --- | --- | --- | --- | --- |
| Gain-framed | NP+ | 3.26 (0.96) | 3.40 | <0.001 |
|  | NP- | -0.31 (0.90) | -0.34 | 0.74 |
|  | NPe | -1.30 (1.07) | -1.22 | 0.22 |
|  | RT | -1.05 (0.10) | -10.40 | <0.001 |
| Loss-framed | NP+ | 2.43 (1.05) | 2.32 | 0.02 |
|  | NP- | -1.70 (0.97) | -1.75 | 0.08 |
|  | NPe | -2.17 (1.32) | -1.64 | 0.10 |
|  | RT | -0.31 (0.09) | -3.54 | <0.001 |
| Self target | NP+ | 4.00 (0.82) | 4.89 | <0.001 |
|  | NP- | -0.62 (0.67) | -0.92 | 0.36 |
|  | NPe | -0.98 (0.96) | -1.01 | 0.31 |
|  | RT | -0.70 (0.10) | -6.97 | <0.001 |
| Charity target | NP+ | 2.41 (0.96) | 2.52 | 0.01 |
|  | NP- | -0.26 (0.57) | -0.45 | 0.65 |
|  | NPe | -1.32 (0.96) | -1.37 | 0.17 |
|  | RT | -0.67 (0.08) | -8.59 | <0.001 |

NP+: Average neural similarity to Neural Profiles associated with the default brain; NP−: Average neural similarity to Neural Profiles associated with the task-engaged brain; NPe: Average neural similarity to Neural Profiles associated with emotion-related terms; RT: Response time

We also investigated whether trial-level neural similarity to cognitive disengagement was predictive of frame-consistent choices within only the subset of self-target trials or only the subset of charity-target trials. Basic Model results within self-and charity-target subsets were qualitatively identical to the results for all trials. That is, for both self-target and charity-target trials, frame-consistent choices were significantly predicted by trial-level neural similarity to the NP+ and response time, but not by trial-level neural similarity to the NP− or NPe (see Table 1).

## Discussion

We show that choices consistent with a typical framing effect best match Neural Profiles associated with the default mode network – not with emotion – while choices inconsistent with the framing effect best match Neural Profiles associated with the task-engaged brain. By combining a large-scale empirical dataset with independent Neural Profiles drawn from meta-analysis, we not only replicated patterns of activations found in previous studies but also systematically tested the relationships of those patterns to cognitive and emotional networks. This approach allowed us to conduct a rigorous neural test of dual-process models of the framing effect.

Our conclusion follows the observation of significant neural activation in PCC and dmPFC for frame-consistent choices, regions associated with the default mode network that are active when the brain is not engaged in performing a task (Gusnard and Raichle, 2001; Hayden et al., 2009). This suggests that frame-consistent choices require limited neural effort and engagement. Note that while we replicate previous findings of amygdala activation during frame-consistent choices (De Martino et al., 2006; Roiser et al., 2009; Xu et al., 2013), we show that neural activity underlying the framing effect extends markedly beyond the amygdala. Furthermore, we note that our Framing Contrast more strongly resembles the Neural Profile of a task-disengaged brain than the Neural Profile of emotional processes, even after accounting for shared variance between Neural Profiles. Finally, we note that at the trial-level, neural similarity to the task-disengaged brain significantly predicts a frame-consistent choice, whereas neural similarity to emotional processes and the interaction of neural similarity to disengagement and emotion do not significantly predict choice.

Our results, coupled with a previous finding that patients with complete bilateral amygdala lesions still exhibit a robust behavioral framing effect (Talmi et al., 2010), indicate that the neural basis of the framing effect is neither specific to the amygdala nor wholly attributable to emotion. While emotions may contribute to the biases seen in the framing effect, our results indicate that susceptibility to the framing effect best reflects varying levels of cognitive engagement during value-based decision-making and does not depend on an interaction between engagement and emotion.

The claim that decision frames drive processes of cognitive engagement is consistent with the observed response time data: choices made during loss frames took longer than choices made during the gain frame, and frame-inconsistent choices took longer than frame-consistent choices. Increased reaction time is often taken as a sign of increased cognitive effort, and though such a reverse inference has been shown to be problematic (Krajbich et al., 2015), our study draws upon additional Neural Profile analyses to substantiate our interpretation. While we cannot rule out non-specific attentional effects (e.g., time-on-task) upon the observed activation in control-related regions (Yarkoni et al., 2009; Brown, 2011; Grinband et al., 2011a, 2011b; Yeung et al., 2011), we note that our fMRI analyses modeled response-time effects by basing the duration of the regressors of interest upon each trial’s response time (Grinband et al., 2011a, 2011b). In this way, we minimized any direct effects of response time upon the reported results and avoided modeling any “mind-wandering” that may occur after participants have reported their choices. Furthermore, we note that our trial-by-trial analyses controlled for response times and still found neural similarity to Neural Profiles associated with the default mode network to significantly predict frame-consistent choices. Thus, neural signatures of cognitive disengagement predict biased choice even after accounting for the significant effect of response times. Finally, given that the degree to which brain activation patterns resemble cognitive disengagement significantly predicts choices within the subsets of gain-framed and loss framed-trials, we can rule out the alternative interpretation that frame itself explains both the brain and behavior pattern.

Our conclusions are bolstered by our sample size (n = 143), which is significantly larger than those of most other neuroimaging studies, including prior studies of the gain/loss framing effect (n = 20 in De Martino et al., 2006; n = 30 in Roiser et al., 2009; n = 25 in Xu et al., 2013). Our large sample provides confidence in our ability to detect true results and making it unlikely that our null results are due to a lack of power (Button et al., 2013). Our conclusions are also strengthened by our use of the large-scale meta-analytic Neurosynth engine (Yarkoni et al., 2011) and our derivation of inferences based on whole-brain networks, not selected clusters of activation (Leech et al., 2011; Gordon et al., 2012; Utevsky et al., 2014; Smith et al., 2015). These methods allowed us to replicate the specific findings of previous studies, to establish a principled test of two competing hypotheses within our new data, and to establish an alternative explanation for framing effects.

We note that the Neural Profiles are created by an automated tool that calculates probabilistic associations rather than deterministic labels, and that they are subject to the biases of how neural activation is interpreted and reported in the fMRI literature (Carter and Huettel, 2013; Chang et al., 2013). However, Neurosynth has been found to be comparable to manual meta-analyses (Yarkoni et al., 2011), and the reporting biases present in Neurosynth are also inherent to all meta-analytic approaches. Therefore we believe that the Neural Profiles of Neurosynth represent the current best synthesis of the fMRI literature’s interpretation of neural activation.

Another limitation of our study involves the lack of simultaneous measurements of specific emotional processes. Though we did not directly measure participants’ self-reported ratings of emotional valence, Neural Profiles for positive and negative emotions exhibited low neural similarity with our Framing Contrast. Future work could build on our findings by obtaining self-report ratings of emotional valence and arousal or trial-to-trial measures of physiological recordings (e.g., skin conductance response). These measures could provide additional insight into the extent to which emotional processes were engaged during (and predictive of) framing-consistent decisions.

Given the overlap between the default mode network and social cognition (Whitfield-Gabrieli et al., 2011; Mars et al., 2012; Jack et al., 2013; Spreng and Andrews-Hanna, 2015), one open consideration is the extent to which our results are due to social cognition rather than minimal cognitive processing. Other studies have found differences in neural framing effects to depend on social context (e.g., receiving feedback from a friend compared to stranger (Sip et al., 2014). In addition, the effects of social feedback on the framing effect have been associated with both the executive control network and the default mode network (Smith et al., 2015). Given that we observed 1) no behavioral differences between our social and nonsocial (charity or self) target conditions, 2) no interaction effects between our target conditions and our gain/loss frame conditions, 3) highly correlated behavioral framing effects between the two target conditions, and 4) trial-level neural similarity to cognitive disengagement to similarly predict choice within the subsets of only self-target and only charity-target trials, it is unlikely that our Framing Contrast results are due to an effect of the “socialness” of the decision target. Instead, it may be that frame-consistent choices and social cognition both evoke default neural networks because both are low-effort cognitive processes.

Our results overturn the previous conceptualization of the framing effect as being solely mediated by emotional, amygdala-driven processes. We show that the framing effect is not uniquely linked to that single region of interest and therefore cannot be wholly attributed to emotional processes. Rather, the biases of the framing effect correspond with differences in neural network activation (e.g., within dACC) that more closely reflect differential levels of cognitive engagement. We note that this finding of differences in engagement does not necessarily imply that dACC and other regions were “disengaged” during the task; on the contrary, activation levels for both Frame-Inconsistent and Frame-Consistent choices were still greater than the implicit baseline of task performance (i.e., the associated regressors had positive signs). Because our event-related task was not designed to cleanly model passive rest, conclusions about absolute levels of engagement cannot be drawn from these data. Future studies could build upon our results using paradigms with well-defined blocks of rest periods.

Though our conclusions are specific to the gain/loss framing effect, future work should determine if they generalize to other supposed examples of Type 1 and Type 2 decision-making biases that have been extensively studied using fMRI, such as loss aversion (Tom et al., 2007; Sokol-Hessner et al., 2012) and impatience in temporal discounting (McClure et al., 2004; Kable and Glimcher, 2007).

“Emotion” versus “reason” has been a popular way to describe the dueling components of Type 1 versus Type 2 decision-making (Kahneman, 2011). Many influential theories of decision-making, however, have accounted for Type 1 decisions without necessarily invoking emotional processes (Simon, 1955; Gigerenzer and Gaissmaier, 2011). Our study, using a large fMRI dataset and an even larger meta-analytic database, suggests that “less cognitive effort” versus “more cognitive effort” is the more accurate characterization of decision-making processes. We show that data from neuroscience can provide novel insights into the processes that underlie well-studied decision science phenomena (Levallois et al., 2012).

## Acknowledgments

The authors would like to thank Anne Harsch and Edward McLaurin for data collection and assistance. The authors would also like to thank Tal Yarkoni for helpful feedback on a previous version of this manuscript. This study was funded by a grant from the National Institutes of Health (NIMH RC1-88680 to S. A. H.) and supported by fellowships from the National Science Foundation (DGF 1106401 to R. L.) and the National Institutes of Health (F32-MH107175 to D. V. S.).

